# Latent effects of winter warming on Olympia oyster reproduction and larval viability

**DOI:** 10.1101/2020.06.01.127977

**Authors:** Laura H Spencer, Erin Horkan, Ryan Crim, Steven B Roberts

**Author notes:** **Corresponding author:** Laura H Spencer, School of Aquatic and Fishery Sciences, Box 355020, Seattle WA 98195.

## Abstract

For ectothermic marine invertebrates living in temperate regions, impacts of ocean warming will vary considerably by season. In many species, reproductive and metabolic processes are tightly linked to the seasonal change from winter to spring, yet we know little about how these processes will shift as winters become milder. This study examined latent effects of winter warming on spring reproduction in the Olympia oyster, *Ostrea lurida*. Adults were collected in autumn from central Puget Sound, WA, USA, and exposed to two winter temperatures (7°C, 10°C) in the presence of food limited (5k algal cells/mL) and food abundant (50k algal cells/mL) environments. Following treatments, adults exposed to elevated winter temperature contained larger oocytes regardless of feeding regime, and those also fed abundant food contained more developed sperm. Adults then spawned in common conditions, and larvae were reared through settlement to assess carryover effects of winter conditions on larval viability. Adults previously exposed to elevated winter temperature (10°C) produced larger larvae, particularly if they were also fed high food levels. More developed gametes and larger larvae suggest that gametogenesis occurred at low levels throughout the winter, possibly resulting in increased maternal provisioning to influence larval size. Interestingly, winter temperature did not impact larval survival, or the timing or magnitude of larval production. In the wild, more developed gametes and larger larvae following milder winters could greatly impact recruitment patterns, possibly benefitting *O. lurida* populations. In the hatchery setting, larval production and survival is not contingent upon winter conditions, and larval survival does not correlate with oocyte and larval size. Our results suggest that *O. lurida* reproduction is resilient to winter warming. Furthermore, as global temperature continues to rise, winter conditions should not be overlooked when examining reproductive cycles of *O. lurida* and other temperate marine invertebrates with similar reproductive cycles.

**Highlights of the manuscript:**

1. Elevated winter temperature resulted in more developed *O. lurida* sperm, larger oocytes, and larger larvae.
2. In experimental settings, *O. lurida* oocyte and larval size upon release did not predict larval survival, but in the wild where rearing conditions are more challenging, winter warming could benefit wild populations by increasing recruitment.
3. Winter temperature did not affect larval production timing or magnitude, indicating that *O. lurida* reproductive capacity is relatively resilient to increasing winter temperatures.

## 1. Introduction

Temperature regulates many reproductive processes in marine invertebrates (Newell & Branch 1980; Hoegh-Guldberg & Pearse 1995). For species that live in temperate regions, reproductive cycles are tightly linked to seasonal temperature changes (Orton 1920). Gametogenesis onset, gamete growth rate, time to ripening, and the act of spawning are all believed to be a function of temperature, along with other environmental drivers such as nutrient availability and photoperiod (Olive 1995; Bates 2005). The precise timing, duration, and frequency of reproductive processes varies among species, but many follow a general seasonal pattern: rapid gametogenesis occurs in the spring; spawning occurs in the late spring and summer; recovery and resorption of residual gametes occurs in autumn, with some early differentiation of next season’s gametes; and reproductive activity slows or ceases in the winter when temperatures drop below a minimum for breeding (Orton 1920; Olive 1995).

As global temperatures rise due to anthropogenic inputs, milder winters are anticipated due to increased sea surface temperature and more frequent marine heat waves (Gentemann et al. 2017; IPCC 2019; Holbrook et al. 2020). Ocean warming will invariably alter marine invertebrate reproductive processes (Edwards & Richardson 2004; Lawrence & Soame 2004). For instance, we know that moderately elevated temperature during gametogenesis and spawning (occurring primarily in spring) can accelerate gamete development (Parker et al. 2018), alter sex ratios (Eagling et al. 2018; Zapata-Restrepo et al. 2019), and increase fertilization rates (Rogers-Bennett et al. 2010; Byrne 2011), and could therefore augment recruitment rates. High summer temperatures can result in adult mortality when spawning coincides with acute heat stress (Mori 1979; Sastry 1966; Samain et al. 2007), reducing the number of breeding individuals or surviving offspring. We know little about how changes to winter conditions will impact reproduction. This oversight could be attributed to the relatively little reproductive activity that occurs in the winter compared to other seasons. There is, however, evidence that winter conditions can carry over to greatly impact physiological processes in other seasons. A 19-year dataset of oyster mortality (*Crassostrea gigas*) and climate variability revealed a lagged response, where high summer mortality occurred following warm winters (Thomas et al. 2018). It is therefore important that we explore more comprehensively how warming during winter months could impact reproduction in temperate ectothermic species.

Many temperate species are thought to enter reproductive diapause in the winter, until temperatures exceed the physiological minimums for breeding in the spring (Orton 1920; Giese 1959; Pearse 1968; Bayne 1976). Winter warming could therefore result in uninterrupted gametogenesis, precocious spawning, or asynchronous sperm release and ovulation, if spermatogenesis and oogenesis respond differently to winter warming (Philippart et al. 2003; Chevillot et al. 2017). Winter warming could impact offspring phenotype by direct impacts to gametes, or indirectly by impacting the physiological processes of progenitors. Egg size, which is associated with nutritional content, typically correlates negatively with maternal environmental temperature (Atkinson et al. 2001; Moran & McAlister 2009; Gosselin et al. 2019). Maternal RNAs and lipid composition, which are utilized during embryogenesis and larval development, could differ if their production or composition are sensitive to winter temperature (Krisher 2013; Leroy et al. 2018). It is also possible that elevated metabolic demand during warmer winters will drain maternal glycogen reserves, resulting in smaller or poor-quality oocytes in the spring (Mathieu & Lubet 1993), particularly if food availability remains limited during winter months (Eppley 1972; Winder & Cloern 2010, but see Testa et al. 2018). Elevated winter temperature therefore has capacity to alter wild populations and cultured stocks through wide-scale shifts in reproductive timing and capacity, and offspring viability.

Impacts of warming on marine species that are of ecological, economic, and cultural importance are particularly concerning. Here, we explore the effects of winter warming in *Ostrea lurida* Carpenter 1864, the Olympia oyster. Once abundant in estuaries along the Northeast Pacific Ocean, overharvest and pollution devastated populations in the early 1900s, and today 2–5% of historic beds remain (Polson & Zacherl 2009; Blake & Bradbury 2012). As the dominant native oyster along the Pacific Coast of North America, *O. lurida* is of cultural and economic significance to tribes and shellfish growers (Peter-Contesse & Peabody 2005; White et al. 2009), is an ecosystem engineer in estuarine habitats (Newell 2004; Coen et al. 2007; Pritchard et al. 2015), and shows some signs of resilience to ocean acidification (Waldbusser et al. 2016; Lawlor & Arellano 2020; Spencer et al. 2020). Restoration efforts are afoot, but *O. lurida* populations are still struggling, and may be further challenged by ocean warming.

*O. lurida* reproduce seasonally, with one or two larval settlement peaks occurring in spring or summer, depending on the location and conditions (Oates 2013; Pritchard et al. 2015). They are hermaphroditic spermcasters, and eggs are internally fertilized then brooded for approximately 10-12 days before being released as D-stage veliger larvae (Coe 1931; Hopkins 1937). Much of the foundational reproductive knowledge for the species was collected in the early 20th century, and the thermal threshold for gametogenesis was then established as ~12.5°C for *O. lurida* in Washington State (the location of our focal population) (Hopkins 1936, 1937). However, recent observations of low-temperature brooding (10.5°C, Barber et al. 2016) and spermatogenesis (~10°C, Spencer et al. 2020) question the concept that Washington State populations cease reproducing below 12.5°C. Winter temperatures in the region typically reach approximately 6.5-8°C (Moore et al. 2008), so even moderately elevated winter temperature could profoundly alter *O. lurida* reproductive phenology by interrupting winter quiescence or altering spawn timing. Sea surface temperature is projected to increase 1.2°C by the 2040’s in the Pacific Northwest (as compared to 1970-1999) under the medium greenhouse gas scenario (Mote & Salathé 2010). However, the region is already experiencing periods of unprecedented warming, such as during the 2013-2015 marine heat wave (coined “The Blob”), which resulted in temperature anomalies exceeding +5°C (Di Lorenzo & Mantua 2016; Gentemann et al. 2017). There were anecdotal reports of poor larval production in an *O. lurida* hatchery in spring 2015 (Ryan Crim, *pers. comm*.), and warm winter temperatures were posited to have interfered with reproduction.

Previously, we explored the combined effects of winter warming and acidification on *O. lurida.* and found that reproduction may begin earlier in the spring following warmer winters, resulting in increased larval production, but when combined with acidification the effects are neutralized (Spencer et al. 2020). Here, we expand the warming-aspect of that study. Adult oysters were exposed to two winter temperatures (7°C, 10°C) in the presence of two feeding regimes (low algal density=5k cells/mL, high algal density= 50k cells/mL). Because winter warming could impact spring reproduction by depleting endogenous energy reserves due to increased metabolic demand (Sokolova et al. 2012), the two feeding regimes were included to assess effects of temperature under an energy limited environment (low algal density) and energy abundant environment (high algal density). The two overwintering temperatures were selected to represent historic (7°C) and elevated (10°C) winter temperatures, with elevated set below the minimum temperature at which *O. lurida* have been found to brood larvae in Washington State (10.5°C in a northern Puget Sound population, Barber et al 2016). Furthermore, the adult oysters were collected from the wild, as opposed to hatchery-reared oysters used in Spencer et al. 2020. Adult growth, survival, and gamete development were monitored during exposure to winter treatments and while spawning in the spring, as well as larval production, size, and survival.

## 2. Methods

### 2.1 Adult winter treatments

Adult *Ostrea lurida* (3.80±0.50 cm) were collected from Mud Bay in Dyes Inlet in Bremerton, WA on November 6, 2017, acclimated to hatchery conditions in filtered (5 um), flow-through seawater and fed live algae *ad libitum*. On December 8th, the adults were divided among eight flow-through tanks (50-L), each with two bags of 50 animals for a total of 100 oysters per tank. Adults were treated in a factorial design to two temperatures (Ambient: 7°C, Warm: 10°C) and two feeding levels (High: 50,000 cells/mL, and Low: 5,000 cells/mL), with two replicate tanks per treatment (200 oysters per treatment in total), for a total of four treatments (7°C+low-food, 10°C+low-food, 7°C+high-food, 10°C+high-food, Figure 1).

**Figure 1:**
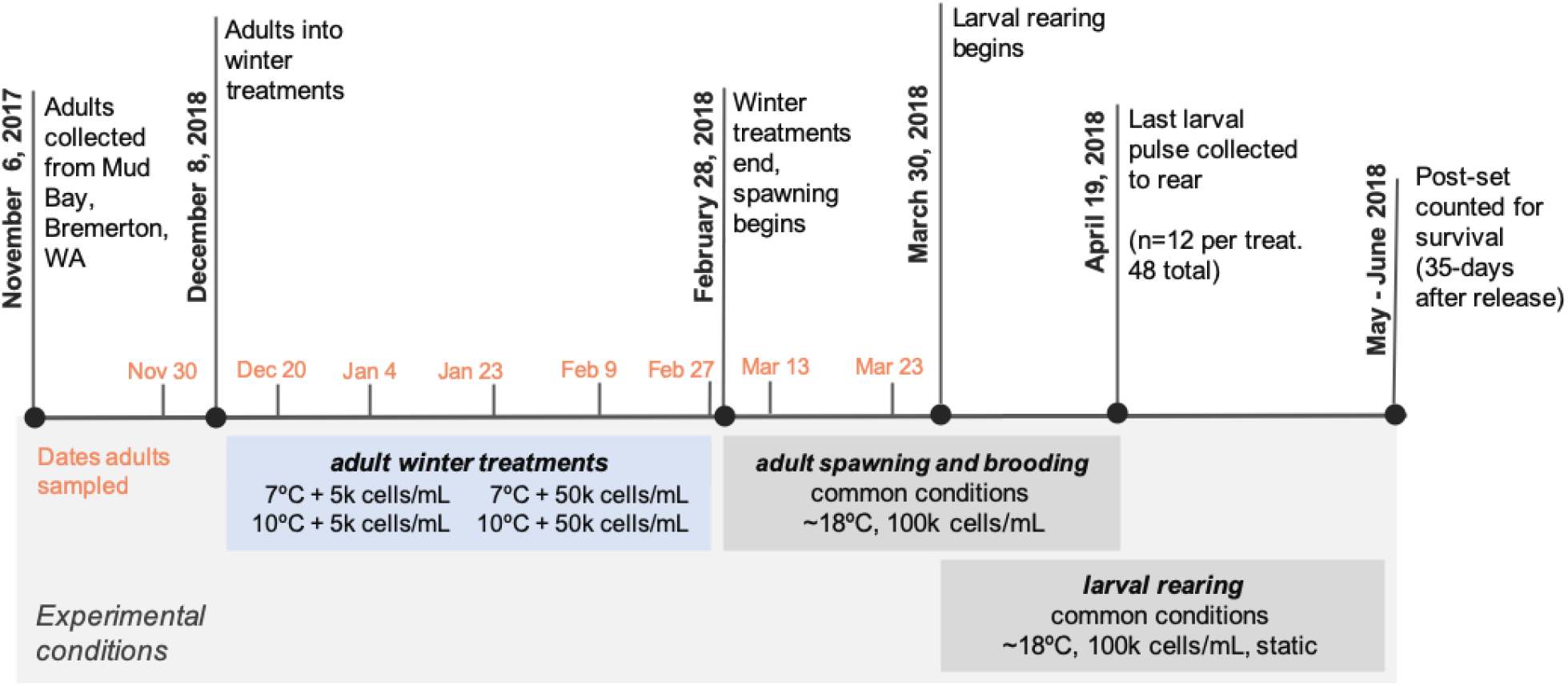
Experimental timeline. Cells/mL indicate the concentrations of live algae given oysters at various stages.

To establish experimental conditions, temperatures were reduced by 0.5°C/day over one and two weeks for the 10°C and 7°C groups, respectively, then maintained for 12 weeks. Temperatures were maintained by recirculating seawater from a reservoir (50-L) through an aquarium chiller (Teco TK-2000 Tank Chiller, 1/3 HP, with built-in heater), before seawater was distributed to experimental tanks at a flow rate of 8L/hr. Feeding rates were maintained using Iwaki metering pumps from a common algae mixture (cocktail of *Tisochrysis lutea, Tetraselmis suecica. Chaetoceros* sp.). Algae cocktail concentration was estimated daily by manual cell counts and adjusting metering pump rates to achieve 50,000 and 5,000 cells/mL for high and low feeding levels, respectively. Temperature was monitored continuously with Avtech temperature probes and recorded with HOBO Pendant Temperature Data Loggers (UA-002-64). Broodstock were cleaned and rotated among experimental tanks and aquarium chillers twice weekly and monitored for mortality. On February 28th the treatment conditions were terminated, and oysters were gradually returned to common conditions for reproductive conditioning and spawning (see section 2.5 for details). Tissue sampling occurred on a regular basis (Figure 1).

### 2.2 Adult tissue sampling

Approximately twice monthly during adult treatments (November 30 - February 27, Figure 1), 10 oysters per treatment were sacrificed to assess growth and gonad development, sampled evenly across treatment replicates and bags within replicates. Adults were also sampled twice during reproductive conditioning (n=12 per treatment, March 13 and 23). Upon sampling, shell height was measured as the distance from hinge to margin, perpendicular to the hinge, and tissue wet weight was estimated by subtracting shell weight from whole wet weight. Gonad tissue was excised by opening the oyster at the umbo, discarding gill tissue, then preserving the whole visceral mass in the PAXgene Tissue FIX system (PreAnalytiX, Hombrechtikon, Switzerland). Fixed tissues were processed for gonad analysis by Histology Consulting Services (Everson, WA).

### 2.3 Gamete development

#### 2.3.1 Gamete stage and sex

The sex and stage of sampled adult oysters were determined from preserved gonad histology sections, using designations adapted from da Silva, Fuentes, and Villalba (2009). As per da Silva, sex was assigned as indeterminate (I), male (M), hermaphroditic primarily-male (HPM), hermaphroditic (H), hermaphroditic primarily-female (HPF), and female (F). Due to the high frequency of hermaphroditism (41.3 % of the 316 sampled oysters), male and female gametes within the same oyster were assigned separate developmental stages, then a dominant gonad stage was assigned for each oyster based on the predominant sex. The da Silva designations were applied for stages 1-3 (1: early gametogenesis; 2: advanced gametogenesis; 3: ripe). Departures from da Silva’s stage 0 (inactive or resting), stage 4 (partially spawned), and stage 5 (fully spawned/resorbing) are as follows: stage 0 in this study represents empty follicles, or no presence of male or female gonad tissue. Stage 4 represents both spawned and resorbing gonad. This method does not include a separate stage 5, due to the very high frequency of residual gametes, and no distinct partially spawned oysters.

Impacts of treatments on gonad development were assessed using Chi-Square or Fisher Exact tests on contingency tables (depending on sample size), which were constructed from counts of gonad sex, male gamete stage, and female gamete stage. Prior to statistical testing, the six sex categories (I, M, HPM, H, HPF, F) were collapsed into four categories by combining HPF and F into one female designation (F), and HPM and M into one male designation (M). Tests were performed for all treatment weeks combined (December 20 - February 27), termination of winter treatments (February 27), and both reproductive conditioning weeks separately and combined (March 13 & 23). To account for multiple comparisons, significance was designated as α=0.01. To determine pairwise differences Fisher Exact post-hoc tests were run using the `pairwiseNominallndependence` function from the rcompanion package (vs. 2.3.7).

#### 2.3.2 Ripe oocyte size

To estimate impacts of winter treatments on maternal provisioning, ripe oocytes were measured upon terminating treatments and during spawning (February 27, March 13 & March 23). The maximum oocyte length was measured for 24 of the largest oocytes in oysters containing stage 3 (ripe) oocytes using a Nikon eclipse Ni microscope and the NIS-Elements BR imaging and measuring software (version 4.60). The maximum length was assessed due to elongated oocyte shape, and the varying orientation of oocytes in mounted histology sections. The number of stage 3 females varied among treatments, and was 6, 14, 7, and 10 for 7°C+low-food, 10°C+low-food, 7°C+high-food, and 10°C+high-food, respectively. Mean oocyte size was calculated for each oyster, then compared among treatments using Two-Way Analysis of Variance.

### 2.4 Larval production

After terminating treatments, adults were spawned to assess impacts of winter treatments on larval production. Following hatchery procedures, continuous, volitional spawning was induced by holding adults in elevated temperature and nutrition in flow-through tanks (20-L at 26-L/hr). Adults from each winter treatment tank were split into two spawning tanks, for a total of four replicate spawning tanks per winter treatment, each with ~25 oysters. Beginning on February 28, temperature was increased 0.5°C/day for 7°C treatments to 10°C (6 days), then all groups increased 1°C/day to 18°C and fed live algae cocktail ad libitum. Tanks were checked daily for veliger larvae, which are released from the maternal brood chamber approximately 10-14 days after fertilization. Once larval release began, larvae were collected daily for four weeks and counts were estimated by hand-counting larvae in triplicate subsamples. Twice weekly tanks were cleaned, adults were inspected for mortality, and then rotated among the tank arrangement.

Larval production timing and magnitude were compared among adult winter temperature and food treatments. Release timing was assessed by comparing the date of onset and date of maximum release using Kruskal-Wallis rank sum tests. Release magnitude was assessed by comparing the total number of larvae released over the 4-week collection period, and the average number of larvae released each day using Two-Way Analysis of Variance and Kruskal-Wallis rank sum tests, comparatively. All metrics were assessed for homogeneity of variance using Bartlett’s Tests for normally distributed data, and Levene’s Test for nonparametric data.

### 2.5 Larval viability

#### 2.5.1 Larval size

To assess impacts of adult winter treatment on larval size, larvae were measured upon release from the maternal brood chamber. After released larvae were counted, excess larvae were preserved directly in −80°C, then measured using a Nikon eclipse Ni microscope and the NIS-Elements BR imaging and measuring software (version 4.60). Mean shell height (distance from hinge to margin, perpendicular to hinge), and mean shell width (longest distance parallel to hinge) were estimated from at minimum 40 larvae per collection from each tank. The number of larval batches that were measured varied among treatments, and was 30, 38, 23, and 29 for 7°C+low-food, 10°C+low-food, 7°C+high-food, and 10°C+high-food, respectively. Mean height and width were compared among parental treatments using lme() from the nlme package to construct the linear mixed-effect models, and Anova() from the car package to construct Analysis of Variance tables and to test for significance using Wald chi-square. Larval preservation method changed part-way through sampling: the first 50 larval samples were sacrificed using ethanol, whereas only fresh water was used to collect later samples. Ethanol use impacted the integrity of preserved larval tissue, resulting in larvae that were on average 5.6 um smaller compared to those that were preserved only with freshwater. We therefore included the use of ethanol as a random variable when testing for effects of parental treatments on larval size (Supplementary Figure 3).

#### 2.5.2 Larval survival

To assess impacts of adult winter treatments on larval survival, a subset of collected larvae were reared through settlement. In total, 48 pulses of larvae were reared, 12 per adult treatment, which were collected over a 19 day period. As multiple females can release larvae on the same day, some larval pulses may represent more than one male x female mating pair, and thus each pulse is henceforth referred to as a “group” (as opposed to “family”). To ensure genetically diverse larval groups within treatments, three groups were collected from each of the four replicate spawning tanks. Upon collecting a larval group, larvae were cleaned of debris using nylon mesh (224 μm) and soaked in fresh water (18°C) for 1 minute. For each larval group, approximately 2,400 larvae were reared in triplicate (800 larvae per tank). Larval stocking error rate (3.1%, mean 824±54 SD larvae) was determined for 12 of the 48 groups by hand counting triplicate samples, taken simultaneous to tank preparation. Larval tanks were constructed from thin-walled polyvinyl chloride pipe (7.6 cm) and nylon mesh (100 μm) placed in individual containers with static water (800±50 mL). Water was changed daily from a common mixture of filtered seawater (<1 μm) and live algae, which consisted of a 1:1 mix by volume of *Chaetoceros muelleri* and *Pavlova pinguis*. for a combined concentration of 100,000 cells/mL. Fourteen days after collecting each larval group, oyster shell fragments (0.5-mL of 224 μm) were sprinkled into larval tanks to serve as settlement substrate. After thirty-five days, survival rate was estimated for each larval tank by hand-counting the number of surviving, metamorphosed larvae, and average post-set survival was calculated from the three replicate tanks for each of the 48 larval groups. Factors influencing larval survival were assessed using quasibinomial generalized linear models (GLM) and Pearson’s Chi-squared tests. Factors tested included adult temperature treatment, adult feeding level, larval size upon release (height, width), the number of larvae released in a group (i.e. approximate brood size), and the date larvae were collected.

## 3. Results

### 3.1 Adult survival and size

During winter treatments there was minimal adult mortality, and no differences in mortality among winter treatments. However, during reproductive conditioning (which began February 26), high mortality occurred in both 7°C groups. Cumulative survival at the end of the experiment was 50% and 70% in the 7°C+high-food and 7°C+low-food groups, and 81% in both 10°C+high-food and 10°C+low-food groups (Figure 2).

**Figure 2:**
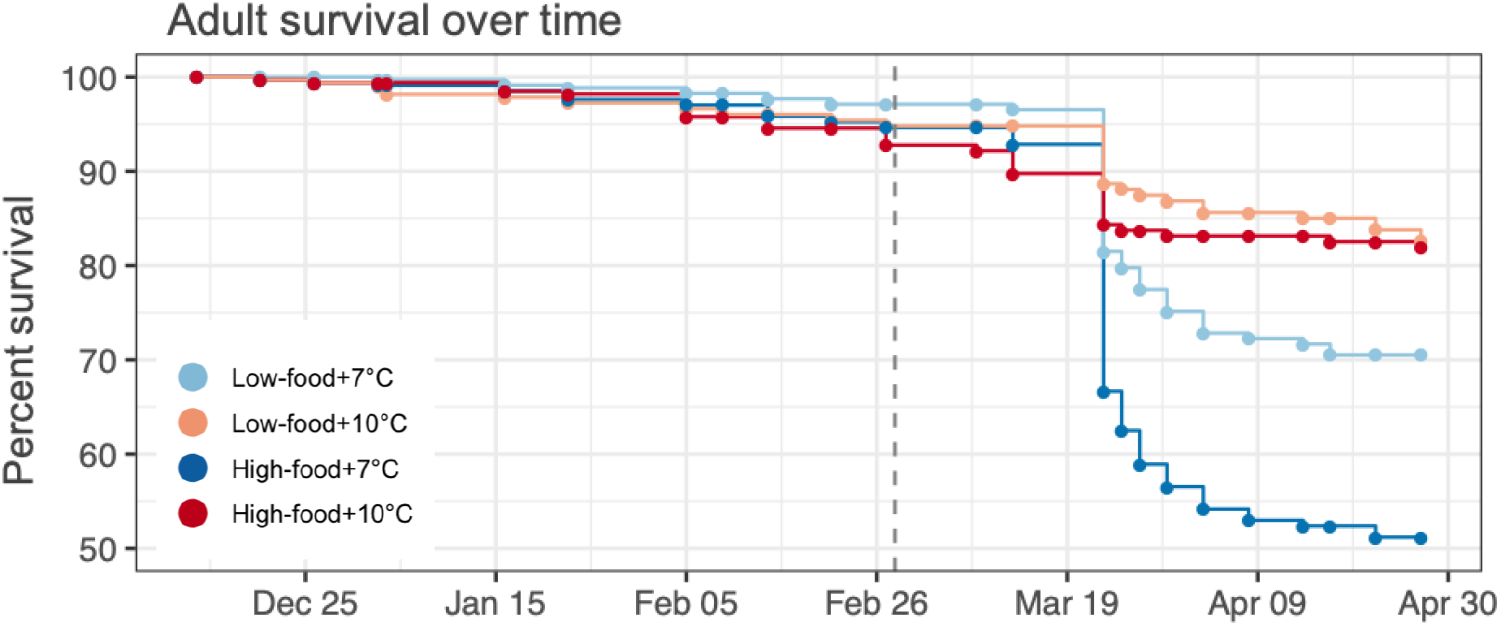
Adult survival over time by winter treatment. There was higher mortality in adults that had previously been exposed to 7°C, particularly those also held in the high food environment, but mortality differences were only observed after they were removed from winter treatments (dashed line) and had entered common spawning conditions.

Adult shell height and wet tissue weight were not affected by winter temperature or feeding level. Wet tissue weight decreased over time in all experimental adults (F(1,280)=11.8, p=7.5e-4) (Supplemental Figure 7). Shell height, which was on average 38.0±5.0 mm, did not change significantly in experimental adults (F(1,294)=3.55, p=0.061).

### 3.2 Gamete development

Winter warming resulted in more developed sperm, but only in the presence of high food (Figure 3, Table 1). There were no significant effects of winter treatments on oocyte stage or gonad sex ratio (Figure 3, Table 1). However, winter warming resulted in larger ripe oocytes (Figure 4). Here we provide more details on sperm development and egg size.

**Figure 3:**
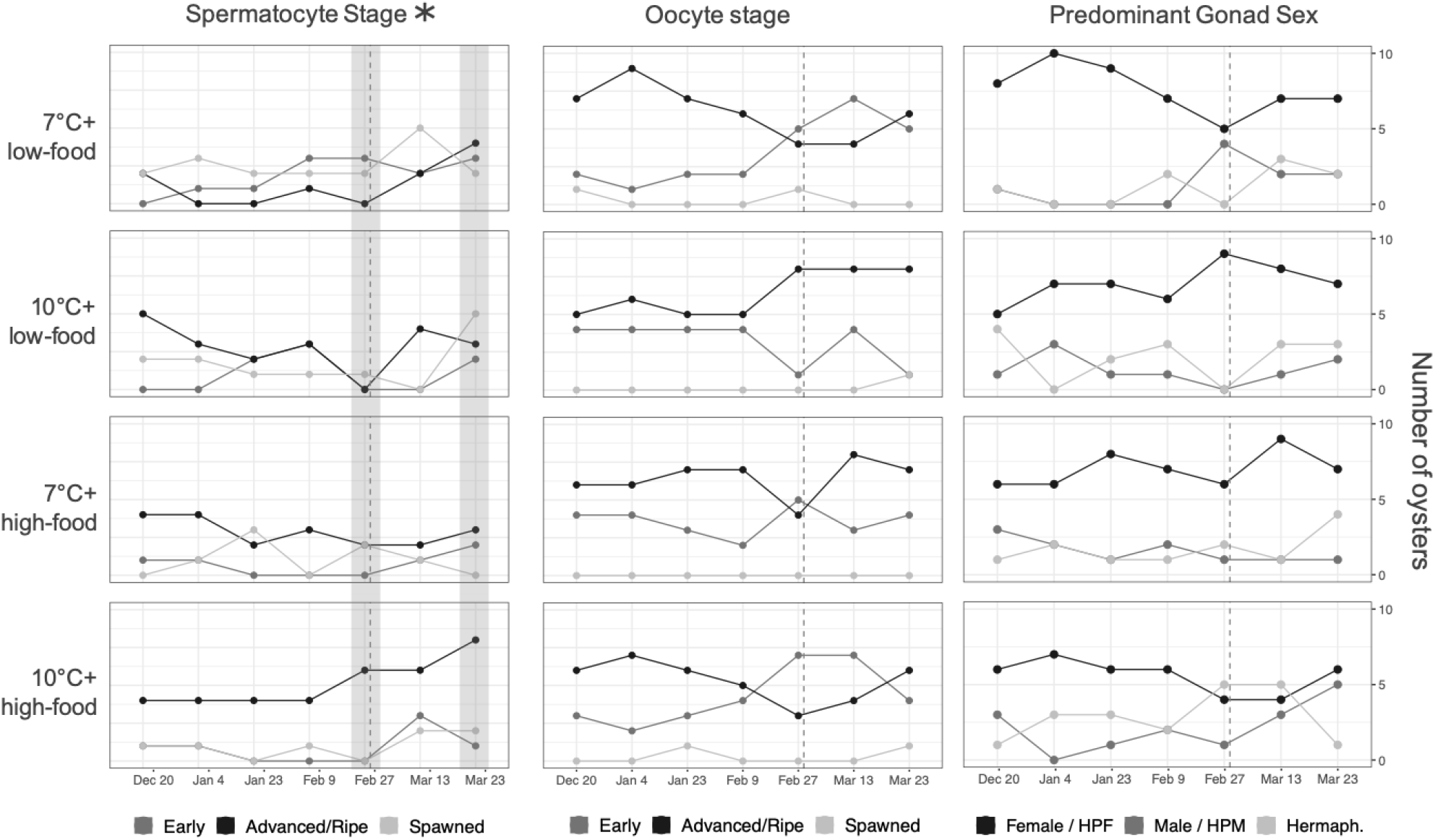
Spermatocyte stage, oocyte stage, and gonad sex for each treatment throughout winter and during spawning. Adults exposed to winter warming and high food had more developed spermatocytes (gray bars) at treatment termination (February 28, dotted line) and while spawning in common conditions (March 23). Oysters were induced to spawn by increasing temperature 1°C/day and feeding with live algae ad libitum. Frequency of Stage 0 oocytes and spermatocytes, indicating none present, are omitted.

**Figure 4:**
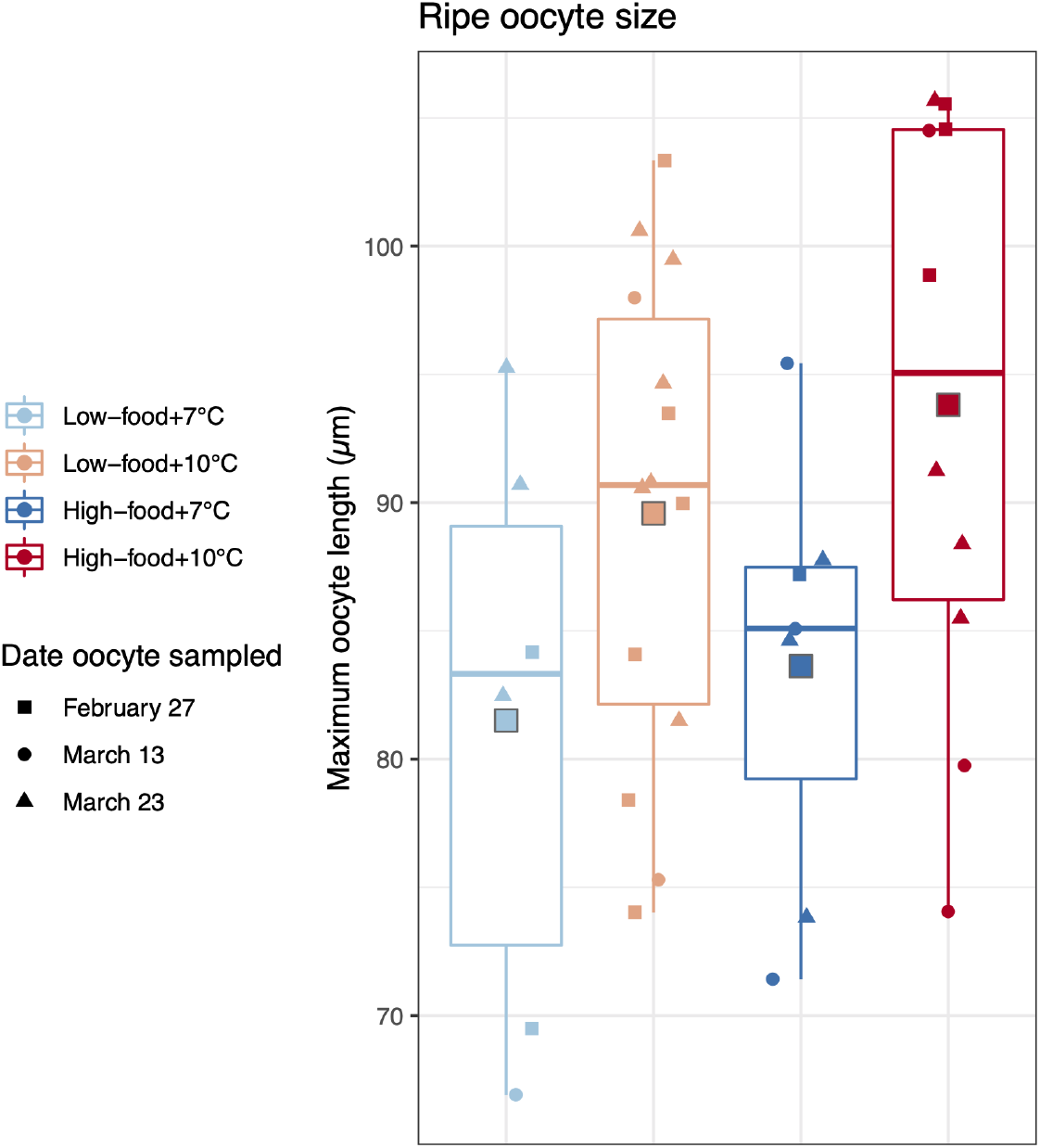
Adults exposed to winter warming (10°C) contained larger ripe oocytes than those held at ambient temperature (7°C), regardless of the feeding regime (F(1,33)=6.06, p=0.019). Each point represents an oyster’s average oocyte length, which we report for all oysters containing ripe oocytes (stage 3) at the termination of treatments (February 27) and during volitional spawning (March 13 & 23). Low-food=5k algal cells/mL, and high-food=50k algal cells/mL. Boxes contain values lying within the interquartile range (IQR), with medians indicated by lines in the middle of boxes. Whiskers extend to the largest value no >1.5*IQR, and points outside the boxes indicate outliers beyond 1.5*IQR. Square points outlined in gray indicate mean oocyte length by adult winter treatment.

**Table 1:** Impacts of treatments on gonad development. Gonad sex, spermatocyte stage, and oocyte stage were compared among winter treatments using all oysters sampled during exposure (Dec 20 - Feb 27), oysters sampled upon termination of exposures (on Feb 27), and those sampled during spawning in common conditions (Mar 13 & 23) by Chi-square contingency table tests, unless otherwise noted. The four treatments included 7°C+low-food, 10°C+low-food, 7°C+high-food, and 10°C+high-food, where high-food was 50k cells/mL and low-food was 5k cells/mL. Where treatment significantly affected gonad development (*), we include pairwise treatment differences according to Fisher Exact tests (due to multiple comparisons, significance was set as α=0.01).

| Date sampled | Spermatocyte Stage | Oocyte Stage | Predominant Gonad Sex (collapsed) |
| --- | --- | --- | --- |
| All adults sampled during treatments (Dec 20 - Feb 27) | $\chi^2 = 25.6$, $p=0.010$ | $\chi^2 = 13.1$, $p=0.38$ | $\chi^2 = 12.4$, $p=0.19$ |
| End of winter treatments (Feb 27) | *$\chi^2 = 25.8$, $p=0.0065$<br>*Pairwise treatment differences:*<br>10°C+high-food vs. 7°C+low-food<br>10°C+high-food vs. 10°C+low-food | $\chi^2 = 16.6$, $p=0.14$ | $\chi^2 = 18.9$, $p=0.020$ |
| 2 weeks in common spawning conditions (Mar 13) | $\chi^2 = 23.4$, $p=0.019$ | $\chi^2 = 8.6$, $p=0.51$ | $\chi^2 = 9.2$, $p=0.43$ |
| 3.5 weeks in common spawning conditions (Mar 23) | *$\chi^2=25.2$, $p=0.0096$<br>*No sign. pairwise differences* | $\chi^2 = 7.7$, $p=0.85$ | $\chi^2 = 8.7$, $p=0.50$ |
| Both spawning dates combined (Mar 13 & 23) | *$\chi^2 = 32.2$, $p=0.0010$<br>*Pairwise treatment differences:*<br>10°C+high-food vs. 7°C+high-food<br>10°C+high-food vs. 10°C+low-food<br>7°C+high-food vs. 7°C+low-food | $\chi^2 = 11.7$, $p=0.49$ | $\chi^2 = 8.7$, $p=0.48$ |

#### 3.2.1 Sperm development

On the final day of adult winter treatments (February 27), sperm developmental stage differed significantly among experimental treatments (*χ*^2^ = 25.8, p=0.0065). More of the 10°C+high-food oysters contained late-stage spermatocytes (60% of male tissue was advanced or ripe) than the 10°C+low-food (0%) and 7°C+low-food oysters (0%) (Figure 3, Table 1). Sperm stage also differed during the spawning phase (March 23: *χ*^2^ = 25.2, p=0.0096; March 13 & 23 combined: *χ*^2^ = 32.2, p=0.0010). The 10°C+high-food group contained more late-stage spermatocytes (58% of male tissue was advanced or ripe) and only 2 oysters fully lacked sperm (8%), compared the other treatments which had a higher proportion without sperm (21%, 42%, and 63% for 7°C+low-food, 10°C+low-food, 7°C+high-food, respectively, Figure 3, Table 1).

#### 3.2.2 Ripe oocyte size

Adults exposed to elevated winter temperature had larger ripe oocytes (stage 3) upon termination of winter treatments and during spawning (Feb 27, Mar 13 & 23 combined) than adults exposed to ambient winter temperature (F(1,33)=6.06, p=0.019, Figure 4). Feeding level had no effect on ripe oocyte size (p=0.32), nor was there an interaction between temperature and food treatments (p=0.77). Mean oocyte length (maximum length) of stage 3 oocytes was 91±14 μm and 83±13 μm for 10°C and 7°C oysters, respectively.

### 3.3 Larval production

Temperature did not influence larval release timing or magnitude as a sole factor. Temperature and food level interacted to influence larval release timing (Food:Temp onset *χ*^2^ = 12.88 p=4.9e-3, Table 2), as did food level as a sole treatment (*χ*^2^ = 10.87 p=9.4e-4). The effect was influenced predominantly by the adults exposed to the 7°C+high-food treatment, which released larvae much later than the other groups (up to one week, Supplemental Figure 2). On average, the total number of larvae released did not differ among treatments (Table 2). However, the total number of larvae released varied less among replicates when adults were fed high levels of food, particularly in those that were also exposed to elevated temperature (Supplemental Figure 2).

**Table 2:** Influence of temperature and food level on larval release timing and magnitude. Kruskal-Wallis Chi-Square Tests were used to assess the number of days until larval release onset. F-statistics from 2-way Analysis of Variances indicate effects of temperature (7°C, 10°C) and food level (5k cells/mL, 50k cells/mL).

| **Metric** | **Temp.** | **Food** | **Temp:Food** |
| --- | --- | --- | --- |
| **Days until larval release onset**<br>*Kruskal-Wallis*<br>*Chi-Square Test* | $\chi^2 = 1.33$<br>p=0.25 | $\chi^2 = 10.87$<br>p=9.8e-4 | $\chi^2 = 12.88$<br>p=0.0049 |
| **Days until peak larval release** | F=1.22<br>p=0.29 | F=0.026<br>p=0.87 | F=4.99<br>p=0.045 |
| **Average no. larvae released each day (~brood size)** | F=0.068<br>p=0.80 | F=0.0008<br>p=0.98 | F=0.0005<br>p=0.99 |
| **Cumulative no. larvae released over the 30-day collection** | F=1.7<br>p=0.22 | F=1.0<br>p=0.33 | F=0.45<br>p=0.52 |

Released larvae were first observed 31 days after entering spawning conditions. Brooded embryos were observed on March 23 during adult oyster sampling, 24 days after beginning the spawning phase. Most brooded larvae were collected from oysters exposed to elevated winter temperature. Of the 48 oysters sampled (12 per treatment), six broods were observed from adults exposed to 10°C+high-food (50% brooding), five from 10°C+low-food (42% brooding), one from 7°C+low-food (8% brooding), and zero in 7°C+high-food.

### 3.4 Larval viability

#### 3.4.1 Larval size at liberation

Adults exposed to elevated temperature produced larger larvae, with larger mean shell width (*χ*^2^=4.69, p=0.030, Figure 5). Adult temperature treatment also interacted with adult food level to affect larval shell width, resulting in significantly larger larvae from adults exposed to 10°C+high-food than those exposed to 7°C+high-food (*χ*^2^=4.28, p=0.039). Adult food treatment alone did not affect larval size (*χ*^2^=2.53, p=0.11).

**Figure 5:**
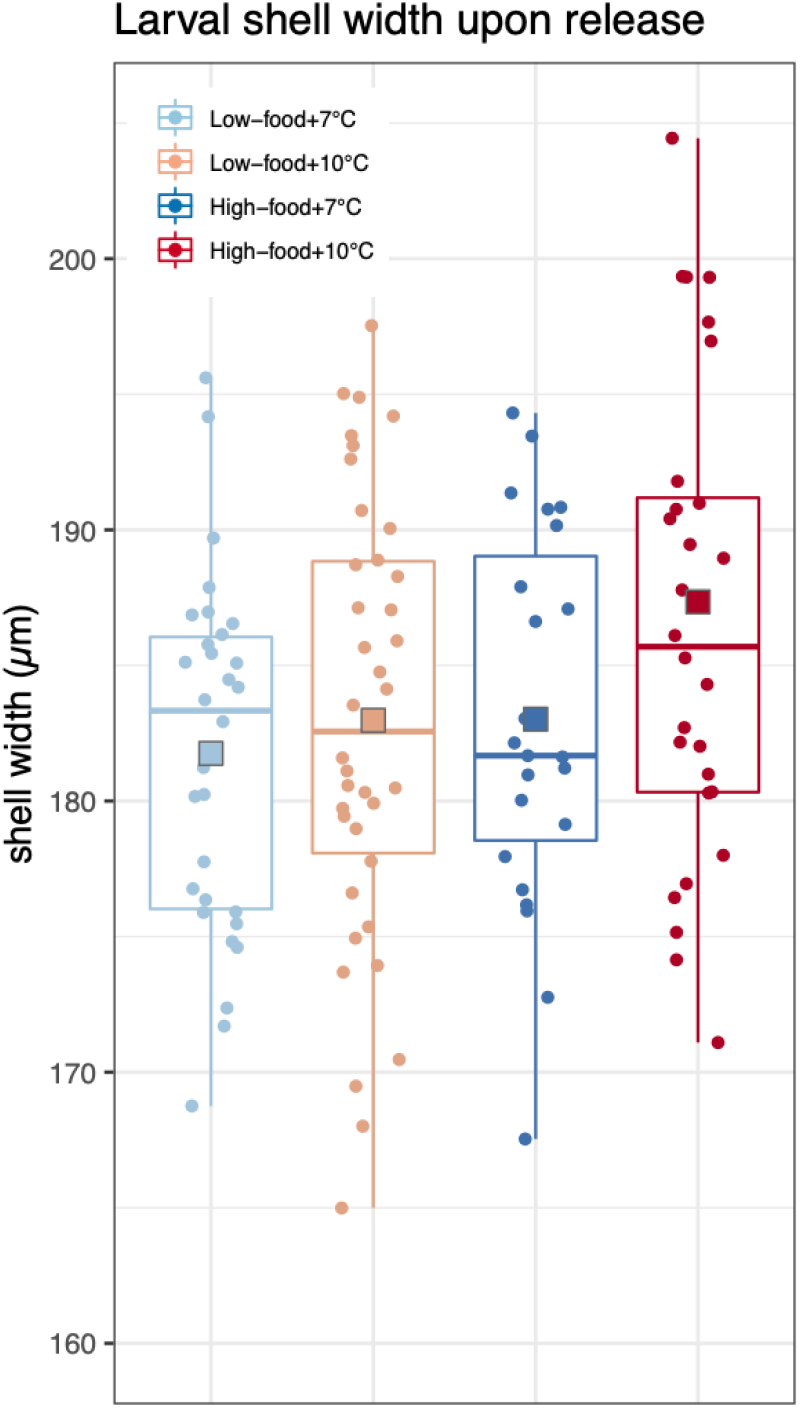
Larvae were larger from adults that had previously been exposed to elevated winter temperature (*χ*^2^=4.69, p=0.030), particularly if they were also fed high food levels (*χ*^2^=4.28, p=0.039). Each point represents the average shell width of a group of veliger larvae released from the maternal brood chamber (N=120). Low-food=5k algal cells/mL, and high-food=50k algal cells/mL. Boxes contain values lying within the interquartile range (IQR), with medians indicated by lines in the middle of boxes. Whiskers extend to the largest value no >1.5*IQR, and points outside the boxes indicate outliers beyond 1.5*IQR. Square points outlined in gray indicate mean larval shell width by parental winter treatment.

#### 3.4.2 Larval survival

Larval survival through metamorphosis (assessed 5 weeks after collection) was not significantly influenced by adult winter temperature (*χ*^2^=0.13, p=0.72) or food treatment (*χ*^2^=0, p=1), nor was there a significant interaction between the two treatments (*χ*^2^=0.84, p=0.36, Figure 6). Larval survival was not associated with larval shell height (*χ*^2^=0.032, p=0.86) or width (*χ*^2^=1.51, p=0.22) upon liberation (Supplemental Figure 4).

**Figure 6:**
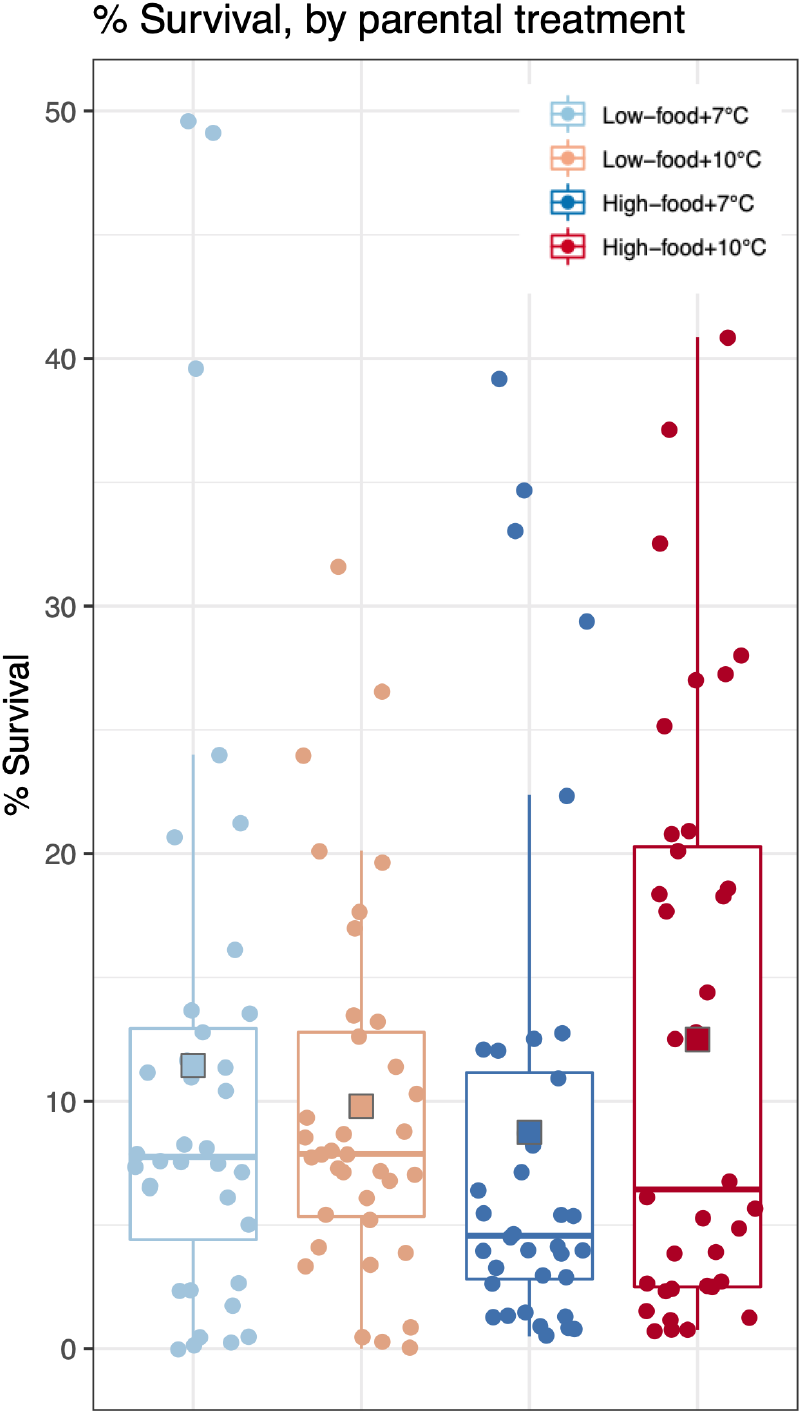
Larval survival (%) by parental winter treatment, which were not associated. For each of the 12 spawning tanks (4 per treatment), three larval groups were reared in triplicate from larval pulses released on different days, for a total of 144 larval tanks (each point = average survival in one larval tank). Low-food=5k algal cells/mL, and high-food=50k algal cells/mL. Boxes contain values lying within the interquartile range (IQR), with medians indicated by lines in the middle of boxes. Whiskers extend to the largest value no >1.5*IQR, and points outside the boxes indicate outliers beyond 1.5*IQR. Square points outlined in gray indicate mean larval survival by parental winter treatment.

All data and code associated with this project are publicly available (Spencer et al. 2021).

## 4. Discussion

This study tested the effects of winter warming on *O. lurida* spring reproduction and larval quality. Adults were exposed to two winter temperatures (7°C, 10°C) and two feeding regimes to examine effects of warming under an energy limited environment (low algal density) and energy abundant environment (high algal density). Winter warming as a sole exposure augmented egg size and larval size, and influenced sperm development in the presence of high-food, but warming did not affect fecundity or larval survival under any feeding regime.

For oysters and many other marine invertebrates living in temperate regions, reproduction is tightly regulated by temperature and seasonality (Korringa 1952; Newell & Branch 1980). Ocean warming in winter could greatly affect these populations via latent effects to gametogenesis and maternal provisioning (among other possible effects) (Ottersen et al. 2001; Lawrence & Soame 2004). In theory, elevated winter temperature increases an ectothermic organism’s metabolic rate (Schulte et al. 2011; Schulte 2015), and, in the absence of increased food, depletes glycogen and lipid reserves, resulting in less energy available for reproduction (i.e. gamete development) and growth (i.e. glycogen/lipid storage, calcification, Sokolova et al. 2012). Therefore, we predicted that elevated winter temperature (+3°C) would result in poor *O. lurida* larval production and/or low quality larvae (i.e. smaller, poor survival), unless they were provided ample exogenous energy (high food). Contrary to expectations, winter warming augmented oocyte and larval size, and had no effect on larval production or survival. We expand on these findings below, and discuss implications for *O. lurida* populations in future ocean conditions.

### 4.1 Gamete development

Winter temperature impacted egg and sperm development. Following exposure to elevated temperature (+3°C), adults contained significantly larger mature oocytes regardless of feeding regime (Figure 4), and more developed sperm when they were also exposed to high food (Figure 3, Table 1). That winter temperature impacted egg size is not unexpected - in ectotherms, egg size often correlates with environmental temperature, both between and within species (Thorson 1950; Atkinson et al. 2001; Fischer et al. 2003; Moran & McAlister 2009). However, the relationship is typically negative, such that elevated temperature results in smaller eggs. We observed the opposite - larger eggs following elevated temperature. We therefore posit that larger oocytes and more developed sperm are both explained by elevated winter temperature triggering or increasing the rate of gametogenesis. While *O. lurida* reproduction was previously thought to cease below 12.5°C for populations along the species’ northern range, recent evidence has indicated that brooding can occur as low as 10.5°C (Barber et al. 2016), and spermatogenesis as low as 10°C (Spencer et al. 2020). Under experimental conditions the observed gamete differences had no bearing on larval production. In a natural setting, however, slight changes in gamete development could greatly influence population dynamics by lengthening the reproductive season (e.g. precocious spawning), and/or increasing spawning rates.

Another possible explanation for the gamete developmental differences is that adults exposed to winter warming experienced less thermal stress when temperature was increased rapidly to induce spawning. Indeed, less mortality was observed in adults that had been exposed to high winter temperature (Figure 1), but only when adults from all treatments were in common conditions and undergoing a 1°C/day increase to induce spawning. The added 3°C increase for the ambient temperature exposed oysters may have been thermally stressful, which could have increased their susceptibility to bacterial infection. This could have resulted in an energetic shift away from vitellogenesis and towards the stress and immune responses in the ambient-temperature group, resulting in high mortality and less developed sperm and smaller eggs (Bayne et al. 1978; Delaporte et al. 2006; Wendling & Wegner 2013; Lokmer & Wegner 2015). While thermal stress might explain the observed gamete differences during spawning, it does not account for developmental differences observed upon termination of winter treatments, prior to spawning (on February 27th, Figure 3, Table 1). Therefore, while thermal stress could have contributed to the observed gamete differences between temperature treatments, it is not likely the sole explanation.

Interestingly, we observed a sex-specific interaction between elevated winter temperature and food level that impacted gamete development. Winter warming increased oocyte size regardless of feeding regime, whereas more developed sperm were only observed following combined warming and high food. From previous studies it is clear that temperature and food availability both likely play significant roles in *O. lurida* reproductive cycles (Coe 1931; Loosanoff & Davis 1952; Bulseco 1982; Joyce et al. 2013). However, few studies have deciphered which environmental factors need to be present to trigger oogenesis and spermatogenesis in *O. lurida*. Our data suggest that temperature may be the dominant factor influencing vitellogenesis, whereas temperature and phytoplankton may both be necessary to trigger spermatogenesis. Foundational studies are still needed to determine the minimum temperature required for vitellogenesis and spermatogenesis in *O. lurida*, and whether supplemental algae is a necessary cue for spermatogenesis, or if it simply accelerates it when combined with elevated temperature.

### 4.2 Larval viability

Adults exposed to winter warming produced larger larvae. Because we also observed larger eggs in those adults (Figure 4), it is very likely that larvae were larger in part because they hatched from larger eggs (Chambers & Leggett 1996). Larvae were measured upon liberation from the maternal brood chamber, which occurs ~10-12 days after fertilization (Coe 1931; Hopkins 1937), therefore size differences may also reflect varying larval growth rates (Helm et al. 1973). Again, larger eggs were observed following elevated winter temperature, which probably reflects increased maternally-provisioned nutrients, and thus more endogenous energy to fuel embryogenesis and pre-feeding larval growth. However, we also observed an interaction between winter temperature and food, such that the largest larvae were produced by adults that were exposed to both elevated winter temperature and high food. Egg size was not similarly affected by food level, therefore egg size alone may not explain the observed larval size differences. The quality or type of biochemical constituents in eggs may have been influenced by winter food treatment, such as fatty acid composition, which correlates closely with larval growth in *Ostrea edulis* (Jonsson et al. 1999) and can be influenced by adult diet (Helm et al. 1991). Adult exposure to winter warming could also have resulted in changes to epigenetic controls of gene expression in larvae, such as changes to DNA methylation, that influenced overall larval physiology and growth rates. DNA methylation patterns can shift upon heat exposure in the Pacific oyster (*Crassostrea gigas*) (Wang et al. 2020), but whether those changes persist and are heritable is not known. To understand the mechanisms by which parental exposure to winter warming influences larval physiology and size, future studies should assay egg biochemical constituents and other plastic, heritable factors (e.g. DNA methylation) alongside egg size and larval performance metrics.

That adults exposed to elevated winter temperature produced larger larvae and eggs is another signal that *O. lurida* may benefit from winter warming. For many marine invertebrates, larger larvae are considered more viable, since they are more capable feeders and swimmers, and can have improved metamorphic success and growth (Marshall & Keough 2007; Marshall et al. 2008). In *Ostrea,* egg size and larval size upon liberation have been positively linked to larval performance. Chilean flat oyster (*Ostrea chilensis*) pediveliger size at release positively correlates with larval growth and spat survival, in addition to egg size and biochemical properties (Wilson et al. 1996), and *O. edulis* larval growth and metamorphic competency positively relates to size and lipid content upon release (Helm et al. 1973; Gonzalez Araya et al. 2012). In other molluscs, larval size is linked to larval metamorphic competency and egg size, such as in the Eastern oyster *Crassostrea virginica*, hard clam *Mercenaria mercenaria*, bay scallop *Argopecten irradians* (Kraeuter et al. 1981; Gallager & Mann 1986), and the Japanese abalone *Haliotis discus hannai* (Fukazawa et al. 2005).

Given the influence of adult winter temperature exposure on egg and larval size in the present study, one would expect to see consistent impacts to larval survival. Interestingly, larval survival was not influenced by parental exposure to elevated winter temperature, nor did survival correlate with larval or oocyte size. This indicates that although winter warming may augment provisioning of gametes and larvae, the effect does not necessarily persist to influence larval competency, at least not under this study’s experimental conditions. While these results contrast previous studies showing an association between size and survival (Kraeuter et al. 1981; Gallager & Mann 1986; Millican & Helm 1994; Wilson et al. 1996; Fukazawa et al. 2005), they align with the related study, Spencer et al. (2020), which also did not detect temperature effects on larval survival. It is probable that the optimal larval rearing conditions in both studies (*e.g*. ample food, regular cleaning) negated the benefits that larger oocytes and larvae can have on larval physiology, and buoyed any larvae with energy deficits. Nevertheless, this and the related study (Spencer et al. 2020) indicate that winter warming does not compromise *O. lurida* larval quality when reared in the hatchery, and larval size when released from the brood chamber is not predictive of survival through settlement. In the wild, where there is higher predation risk, conditions are more stressful, and phytoplankton abundance is less consistent, winter warming may benefit Puget Sound *O. lurida* populations by increasing larval recruitment due to increased size and/or growth rate (Swezey et al. 2020).

### 4.4 Larval production

Larval production during the 30-day collection period was unaffected by winter temperature, regardless of algal density. The results are in contrast with a complementary study, Spencer et al. (2020), in which elevated temperature exposure prior to spawning resulted in more larvae. The present study specifically expands Spencer et al. (2020), to use a new *O. lurida* population collected from the wild rather than oysters that were bred in captivity. The response of *O. lurida* reproduction to elevated winter temperature may be conditional upon gonad stage prior to treatment, and population-specific reproductive traits (Barber et al. 1991; Silliman et al. 2018). For instance, in this study the percentage of female broodstock was unusually high upon collection (63%), and many already contained late-stage oocytes (55% of females). In comparison, only 28% of all oysters sampled in Spencer et al. (2020) were female, 33% of which contained late-stage oocytes. Oysters that enter the winter with late-stage oocytes may be less influenced by winter conditions, and may require less time and energy for maturation in the spring. The developmental status of Olympia oyster oocytes entering the winter season may therefore influence how winter temperature affects spring reproduction. It must be noted that when we first observed brooded larvae in sampled adults (March 23rd), we found that more of the high-temperature exposed oysters were brooding (42%-50%, vs. only 0%-8% of the low-temperature adults). Differing brooding frequencies could be a signal that winter warming did indeed result in precocious spawning by functional females, however this limited data does not allow us to draw any concrete conclusions. Ultimately, that larval production was unaffected by overwintering treatments suggests that *O. lurida* are capable of withstanding a range of winter conditions without compromising spring larval production. Furthermore, this and Spencer et al. 2020 studies find no evidence that the poor larval production in the *O. lurida* hatchery (Ryan Crim, *pers. comm*.) was attributed to the 2013-2015 marine heat wave (Gentemann et al. 2017).

### 4.5 No carryover effects of winter feeding regime

Parental feeding regime was not the focal treatment in this study, therefore we will not expound its effects in detail. However, it is interesting that adult winter food level did not influence gametes or larvae as a sole factor. Prior work in *Ostrea* spp. has indicated that parental nutrition at various phases can greatly influence the size, growth rate, and survival of offspring (Lannan et al. 1980; Millican & Helm 1994; Wilson et al. 1996; Berntsson et al. 1997; Gonzalez Araya et al. 2012; Marshall & Keough 2007). Our results indicate that for *O. lurida* to successfully reproduce in the spring, nutritional needs during the winter are flexible. In the wild, warming-induced changes to winter phytoplankton availability or composition (Cavole et al. 2016; McCabe et al. 2016; Peterson et al. 2017) may not have pronounced effects on spring reproduction. Of course, our experimental conditions must be considered, such as the high quality larval diet, which could have offset any nutritive differences that adult winter diet imparted to offspring. Additionally, adult diet was only limited during treatment, not during spawning, which could have obfuscated effects of winter malnourishment. The adults tested may also have been buffered by high amounts of endogenous energy reserves prior to entering treatments. Starving adults of various condition index through the spawning and brooding phases would expose whether *O. lurida* rely solely on endogenous glycogen to provision gametes (“capital breeder”), or exploit and require exogenous energy for gametogenesis (“income breeder”)(Bayne 2017).

## 5. Conclusion

There is growing evidence that *O. lurida* is more equipped than other bivalves to handle shifting ocean conditions (Waldbusser et al. 2016; Gray et al. 2019; Lawlor & Arellano 2020; Spencer et al. 2020). While we recognize the limitations of our hatchery-based experimental design in predicting how wild *O. lurida* will respond to ocean warming, our findings do suggest that *O. lurida* reproduction and larval viability are, at the least, not highly sensitive to winter warming, and at best may benefit from it. Our results also provide more evidence that *O. lurida* reproduction is not “on pause” all winter (Barber et al. 2016; Spencer et al. 2020), but rather gametogenesis is likely occurring at temperatures lower than the previously established threshold for reproduction in Washington State populations (12.5°C, Coe 1931, Hopkins 1936). Winter conditions should therefore not be overlooked when examining reproductive cycles in *O. lurida* and other temperate oysters, particularly as the oceans continue to warm and marine heat waves occur more frequently. Additionally, based on histological samples collected throughout the winter, we think that many residual gametes remained viable throughout the winter in our adult oysters. Therefore, winter warming could have influenced larval size by altering factors other than size or stage in gametes, such as macromolecules or RNAs deposited in eggs, or epigenetic modifications to sperm or eggs (e.g. DNA methylation changes). Examining biochemical composition, gene expression, and/or epigenetic factors in gametes after warm and cool winters could reveal the mechanisms responsible for the carryover effects observed here. There are, however, many gaps in our understanding of *Ostrea* spp. reproductive systems, which constrain interpretation of complex molecular data collected from *O. lurida*. It is therefore critical that we untangle how precisely environmental drivers (e.g. temperature, food, tidal cycle, allosperm), and internal drivers (e.g. age, source population), control reproduction across seasons, and in the context of ocean warming. Given the potential resilience of *O. lurida* to ocean change, its reproductive processes may become more pertinent to those culturing, harvesting, conserving, and restoring marine invertebrates.

## 6. Acknowledgements

Our gratitude to the following people who assisted with this project: the NOAA Manchester Research Center and Puget Sound Restoration Fund (PSRF) provided facilities and materials, and PSRF staff members Stuart Ryan, Morgan Adkisson, and Josh Bouma assisted with daily feeding and maintenance. Duncan Greeley and Anne Spencer helped with the larval husbandry; Kaitlyn Mitchell assisted with larval measurements; committee members Jackie Padilla-Gamñio, Rick Goetz, and Jennifer Ruesink advised and supported this project.

This work was supported in part by the National Science Foundation Graduate Research Fellowship Program. This publication is partially funded by the Joint Institute for the Study of the Atmosphere and Ocean (JISAO) under NOAA Cooperative Agreement NA15OAR4320063, Contribution No. 2020-1078.

